# Antimicrobial resistance genes predict plasmid generalism and network structure in wastewater

**DOI:** 10.1101/2022.05.11.491481

**Authors:** Alice Risely, Thibault Stalder, Benno I. Simmons, Eva M. Top, Angus Buckling, Dirk Sanders

## Abstract

Plasmids are mobile genetic elements that can act as mutualists or parasites to their bacterial hosts depending on their accessory genes and environment. Ecological network theory predicts that mutualists, such as plasmids with antimicrobial resistance (AMR) genes in the presence of antimicrobials, should act as generalists, while plasmids without beneficial genes are expected to be more specialised. Therefore, whether the relationship between plasmid and host is mutualistic or antagonistic is likely to have a strong impact on the formation of interaction network structures and the spread of AMR genes across microbial networks. Here we re-analyse Hi-C metagenome data from wastewater samples and identify plasmid signatures with machine learning to generate a natural host-plasmid network. We found that AMR-carrying plasmids indeed interacted with more hosts than non-AMR plasmids (on average 14 versus 3, respectively). The AMR plasmid-host subnetwork showed a much higher connectedness and nestedness than the subnetwork associated with non-AMR plasmids. The overall network was clustered around Proteobacteria and AMR-carrying plasmids giving them a crucial role in network connectivity. Therefore, by forming mutualistic networks with their hosts, beneficial AMR plasmids lead to more connected network structures that ultimately share a larger gene pool of AMR genes across the network.

## Introduction

Plasmids play a key role in the spread of antimicrobial resistance (AMR) and other genes (e.g., metal resistance, biodegradation, virulence), both within and between bacterial taxa (Bennett 2008; Martínez 2008; Dang *et al*. 2017; San Millan 2018; Acman *et al*. 2020). Understanding the ecological mechanisms that underpin plasmid transmission within bacterial communities is important for combating the spread of AMR and associated bacterial epidemics (Dimitriu *et al*. 2021). However, our knowledge about plasmid-host interactions is mostly gained from laboratory research on a limited number of bacteria and plasmids. Therefore, there remains considerable uncertainty surrounding the role of plasmids within larger communities and the resulting plasmid-host networks in nature. This limits our understanding of the ecological and evolutionary processes driving plasmid transmission across natural microbial communities.

Ecological interaction network analysis can provide crucial insights into community structure (Kaiser-Bunbury *et al*. 2017), the speed of disease transmission (González-Salazar & Stephens 2012), the dynamics of coevolution (Guimarães *et al*. 2017) and community stability (Thébault & Fontaine 2010; Veron *et al*. 2018). Recently, the ecological network approach has been applied to microbiological systems to elucidate the ecological mechanisms that underpin microbial dynamics (Flores *et al*. 2013; Weitz *et al*. 2013; Coyte *et al*. 2015; Wang *et al*. 2016). The structure of ecological interaction networks is often related to the prevalent type of interaction between species, with antagonistic and mutualistic interactions associated with different network structures (Thébault & Fontaine 2010; Montesinos-Navarro *et al*. 2017). Theory predicts that the structure for each network type leads to increased stability: mutualistic networks tend to be nested with generalists linking the whole network while antagonistic networks often have a modular structure dominated by specialist interactions (Thébault & Fontaine 2010). Observation studies support theoretical predictions that ecological interaction networks dominated by antagonists tend to have fewer generalist interactions than mutualistic networks (Fontaine *et al*. 2009; Thébault & Fontaine 2010; Montesinos-Navarro *et al*. 2017; Newbury *et al*. in press). Network structure is also influenced by the coevolutionary history of interacting species, because many interactions are evolutionary constrained (Segar *et al*. 2020). For example, due to coevolutionary arms races we can expect interactions in antagonistic networks to be characterised by stronger phylogenetic signal than in mutualist networks (Rohr & Bascompte 2014).

These theories can potentially be applied to bacteria-plasmid networks, because plasmids have highly variable host ranges (Suzuki *et al*. 2010; Klümper *et al*. 2015) and can be parasitic and mutualistic to their bacterial hosts (Lili *et al*. 2007; Harrison & Brockhurst 2012). In many cases, plasmids simply impose a fitness cost on their host, and survive in host communities through the evolution of a lower fitness cost, high transmission rates, high fidelity of partitioning, and sophisticated killing systems ensuring their stable presence within bacterial lines (Jensen & Gerdes 1995). However, plasmids often carry accessory genes, such as AMR genes, that promote the survival of the host bacteria under certain environmental contexts (Lili *et al*. 2007; Harrison & Brockhurst 2012). Plasmids that carry context-dependent beneficial genes may therefore be expected to interact with more bacterial taxa than those that don’t (Fig. 1A-C). This will be driven by both ecological dynamics (selection for hosts associated with a mutualistic plasmid) and coevolutionary dynamics between hosts and plasmids leading to the evolution of generalism.

**Figure 1.**
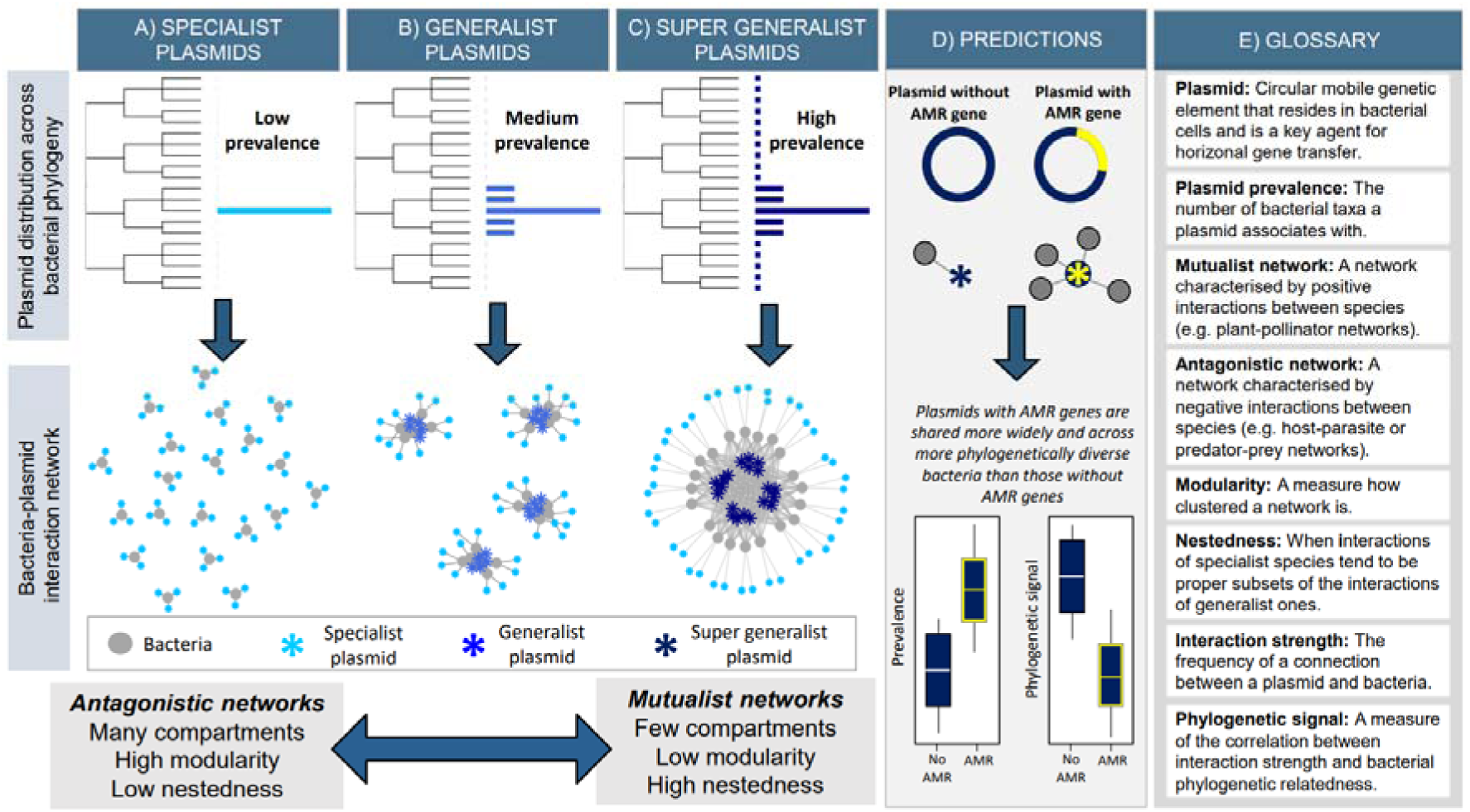
Predictions for plasmid-host network structure. A-C) Three different hypothetical network structures based on 20 bacterial taxa and 60 plasmids, assuming some level of phylogenetic signal in interaction strength. In each network, 40 out of the 60 plasmids are highly specialist (light blue) and interact with only one bacterial taxa. Network structure changes dramatically if the remaining twenty plasmids are A) also specialist; B) generalist but limited to specific clades (royal blue); or C) super generalist (navy blue) with weak interactions across other clades. Ecological theory suggests that antagonistic networks tend to be made up of high number of specialists that lead to networks with many separate compartments, high in modularity, and low in nestedness. On the contrary, mutualist networks are structured by generalists and form networks that have few compartments, and low in modularity, and high in nestedness. D) Because antimicrobial resistance (AMR) genes are likely to have beneficial effects on bacterial fitness within the context of a wastewater community, we predict that plasmids that carry AMR genes are likely to have higher prevalence and lower phylogenetic signal than plasmids without AMR genes. Because increases in prevalence of at least some plasmids lead to more connected networks (A-C), plasmids with AMR should lead to more connected networks. E) Glossary of terms used throughout the article.

Over 10,000 plasmids have been described and referenced to date (likely only a small fraction of real numbers), the majority of which are associated with the bacterial phylum Proteobacteria, especially Alpha- and Gammaproteobacteria (Redondo-Salvo *et al*. 2020). Both specialist and generalist plasmids have been identified, yet an analysis of a subset of described plasmids suggests that up to 60% of plasmids are associated with multiple bacterial host species, and that transmission is limited by host phylogeny with only a small number of super-generalist plasmids (Suzuki *et al*. 2010; Redondo-Salvo *et al*. 2020). However, the link between a plasmid’s range of hosts in a natural microbial community and its likely effect on bacterial host fitness in that community remains unknown. Note that in this study we distinguish between the concepts of host range and generalism. Host range, as it has been defined by plasmid biologists, is the ability of a plasmid to replicate in phylogenetically diverse hosts (i.e., narrow host range versus broad host range). This view of plasmid host range does not necessarily reflect the actual spread of the plasmid in a bacterial community (e.g., a broad-host-range plasmid could be found in only one phylogenetically distinct species of a community and thus be considered a specialist while a narrow-host range plasmid could be found in many different closely related species and be a generalist plasmid). Hereafter we refer to generalist plasmids as plasmids that are found in many species within a community, and specialist plasmids as those that are found in very few host species.

To understand the functional role of plasmids in ecological communities, ranging from soil, to wastewater, to the gut microbiome, we need to identify interactions between plasmids and their hosts in their natural environment. Recently, proximity-ligation methods such as Hi-C allow us to overcome previous sampling limitations and have been used to detect associations between DNA molecules originating in the same cell within microbial communities (Stalder *et al*. 2019; Kent *et al*. 2020; Yaffe & Relman 2020). Moreover, machine learning techniques that can identify plasmids based on their genomic signatures, such as GC content and (tetra)nucleotide composition-specific sequences, with 96% accuracy, allowing for the detection of plasmid genomes that have not officially been described (Krawczyk *et al*. 2018; Pellow *et al*. 2020). Together, these novel methods allow us to quantify host range distributions of plasmids in natural bacterial communities.

Here we apply an ecological network analysis to test the effect of mutualistic interactions on the structure of a natural plasmid-bacteria network, using plasmid-bacteria interaction data from Stalder *et al*. (2019) that used Hi-C to link mobile genetic elements to their bacterial hosts in a wastewater sample. Here we advance/expand on that study by using machine learning methods to identify clusters of plasmid contigs that original from the same cell (hereafter termed ‘putative plasmids’), and test whether AMR presence on these putative plasmids is associated with altered interaction distributions with 374 bacterial metagenome-assembled genomes (MAGs).

While plasmids can carry a range of host-beneficial genes, we make the assumption that putative plasmids that carry AMR genes will be on average more beneficial than those that don’t, because the concentrations of antibiotics in wastewater are likely to confer an advantage to at least some AMR genes (Karkman *et al*. 2018). In this context, we predict that plasmids that carry AMR should interact with more bacterial hosts (i.e., have higher ‘prevalence’; Fig. 1D, see Fig. 1E for glossary of terms) than plasmids without AMR, and thereby lead to more connected and generalist networks. Because evolutionary arms races in antagonistic networks tend to generate strong phylogenetic signal (Rohr & Bascompte 2014), we would also expect that AMR-carrying plasmids to demonstrate weaker phylogenetic signal than those without AMR genes (Fig. 1D). We further discuss the potential of network structures in promoting the spread of AMR.

## Results

The inferred wastewater bacteria-plasmid network was made up of 374 bacterial MAGs (metagenome-assembled genomes) and 289 putative plasmids. Most MAGs identified belonged to either the class Betaproteobacteria (Phylum Proteobacteria), Gammaproteobacteria (Phylum Proteobacteria), Clostridia (Phylum Firmicutes), Bacteroidia (Phylum Bacteroidetes) or Actinobacteria (Phylum Actinobacteria) (Fig. 2A). Bacterial MAGs associated with an average of 3.5 putative plasmids (median = 1, min. = 0, max. = 47), whilst putative plasmids associated with an average of 4.5 MAGs (median = 2, min. = 1, max. = 80). Note that this does not equate to one bacterial cell having 47 plasmids; rather, 47 putative plasmids were found to be associated with that MAG across its entire population within the wastewater community. While MAG relate to a metagenomic bin that can be asserted to be a close representation to an actual individual genome, here the quality of most of the MAG identified (according to genome size and completeness scores computed by CheckM) does not allow us to do this assumption and rather consider the MAGs not as one bacterial cell’s genome, but the genome (or fragment of a genome) of closely related strains within a species.

**Figure 2.**
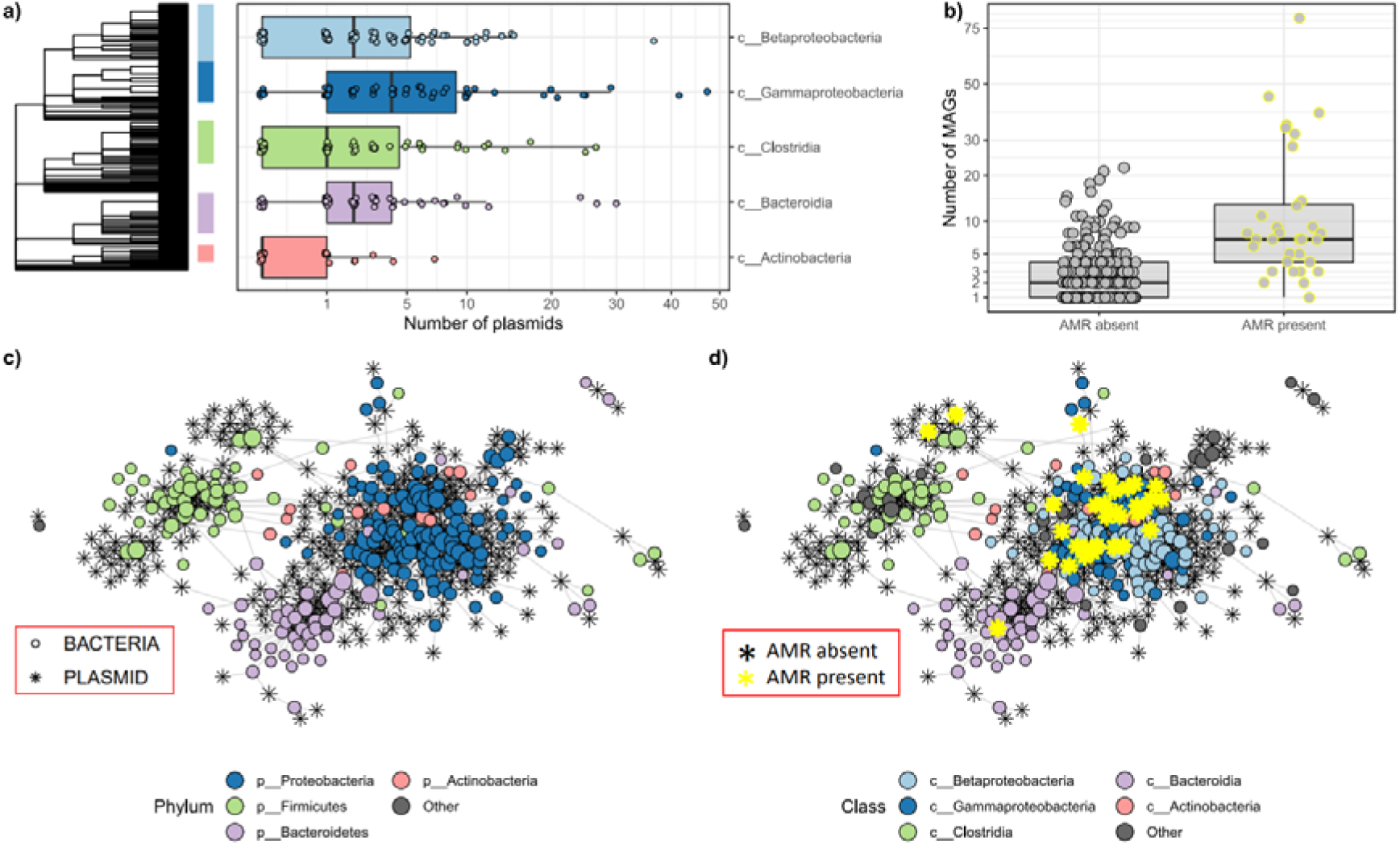
MAG-putative plasmid networks based on normalised Hi-C linkage. A) Distribution of number of plasmids per MAG, split by bacterial class; B) Number of MAG associations per putative plasmid, split by whether plasmids were associated with antimicrobial resistance genes (AMR); C) The full MAG-putative plasmid network made up of 374 MAGs and 289 putative plasmids, with MAGs coloured by phylum and plasmids represented by stars; D) Same network as C) but coloured by class and the position of AMR plasmids highlighted in yellow. Nodes are sized by their network degree.

MAGs that associated with a high number of plasmids were distributed across the phylogenetic tree, although MAGs belonging to Betaproteobacteria, and Gammaproteobacteria tended to associate with a higher number of plasmids (Fig. 2A). Putative plasmids that were associated with AMR genes tended to be more widely distributed across MAGs than those that were not associated with AMR genes (mean AMR = 14, mean no AMR = 3; Wilcoxon W = 1379, p < 0.0001; Fig. 2B). The full network clustered strongly by bacterial taxonomy, with MAGs belonging to Proteobacteria, Firmicutes, and Bacteroidetes largely clustering separately (Fig. 2C). The large majority of AMR plasmids clustered together with Proteobacteria, with classes Betaproteobacteria and Gammaproteobacteria clustering together and sharing most of the AMR plasmids (Fig. 2D).

We next investigated the role of AMR genes on network structure by comparing sub-networks based on whether putative plasmids associated with AMR or not (Fig. 3). When AMR putative plasmids were excluded (Fig. 3A), networks were more modular (higher number of not linked sub networks) and less nested (see Figure 1E) than networks based on solely AMR putative plasmids (Fig. 3B). Putative plasmids carrying AMR genes further connected Proteobacteria to other phyla linking large parts of the whole network.

**Figure 3.**
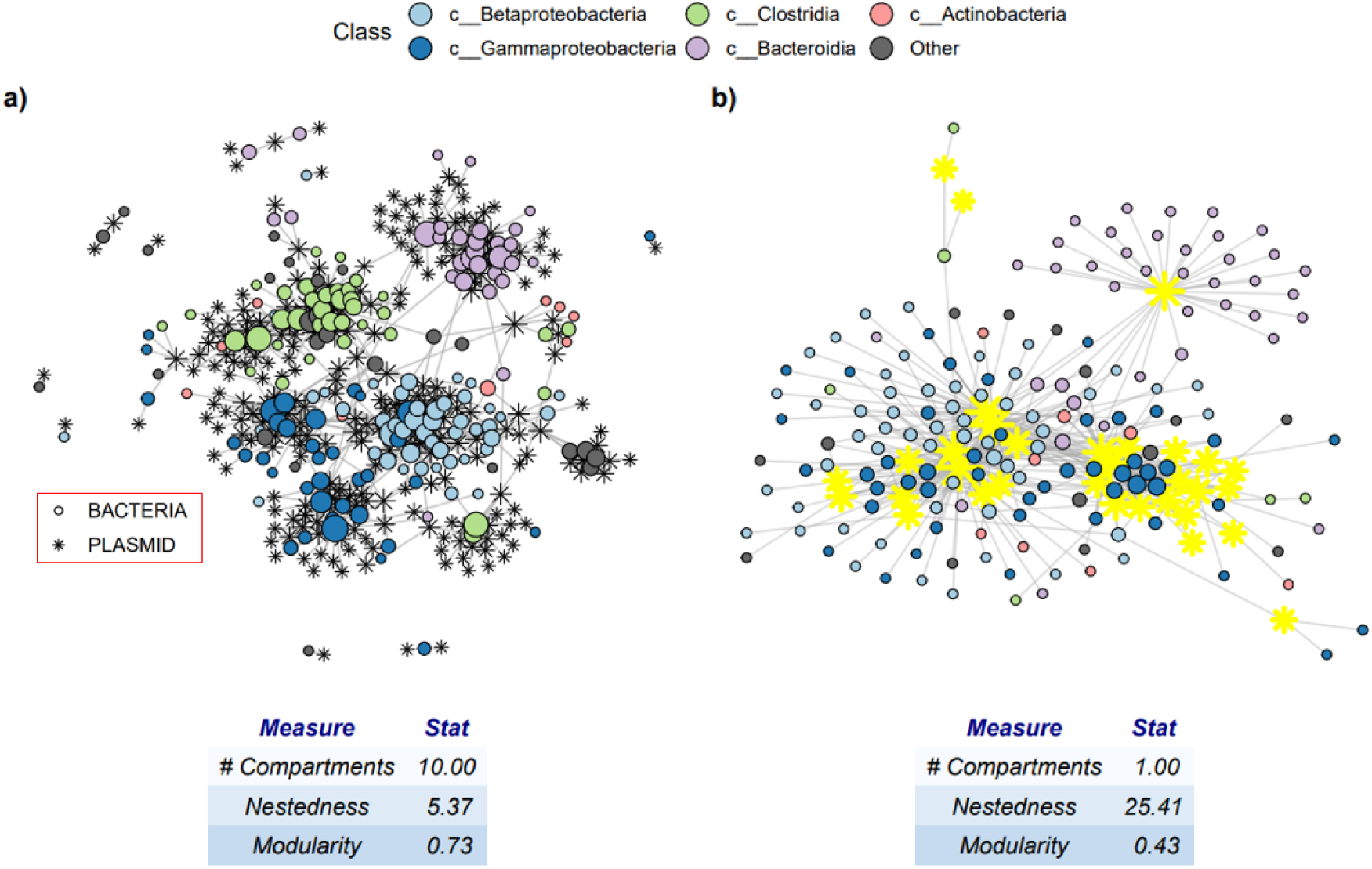
AMR genes and network structure. Sub-networks and network statistics representing MAG Hi-C associations with A) plasmids without AMR and B) plasmid with AMR. Stars represent plasmids and circles MAGs, with AMR plasmids highlighted in yellow. Nodes are sized by their network degree.

We next visualized the distribution of the 15 most prevalent putative plasmids (super generalists) across the bacterial phylogenetic tree (Fig. 4). The most prevalent putative plasmids were largely shared amongst members of the same phyla, although some were occasionally shared more widely. Putative plasmids that were associated with Proteobacteria were often shared across both Beta- and Gamma-proteobacteria (Fig. 4). Whilst most putative plasmids remain undescribed, plasmid 3 (Fig. 4) was identified as a broad range plasmid belonging to the IncP-β group.

**Figure 4.**
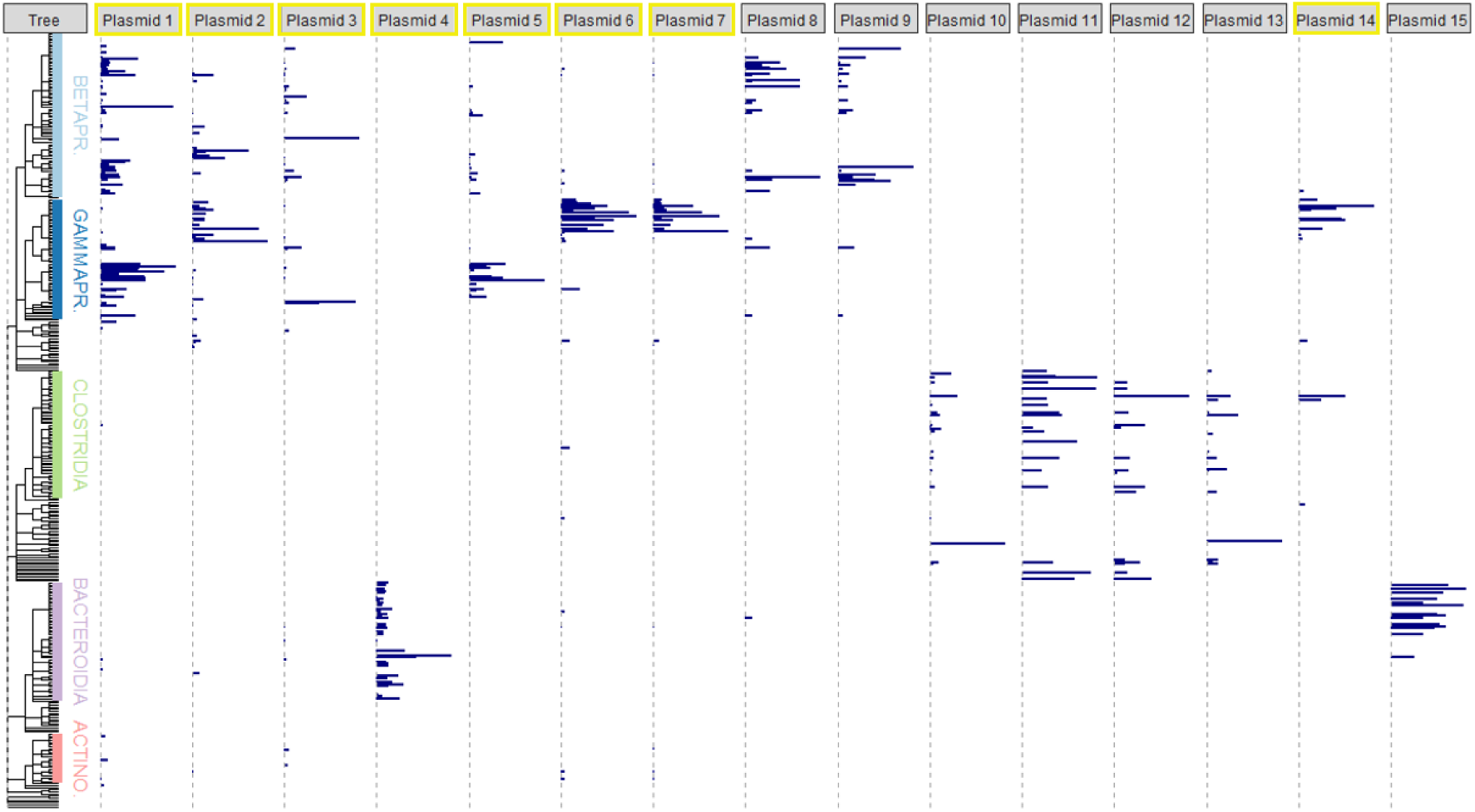
Phylogenetic distributions of the most prevalent plasmids across 374 MAGs. Phylogenetic weighted distributions of the 15 most prevalent putative plasmids (i.e. plasmids with associations to the highest number of MAGs), ordered by how many MAGs they associate with. Bar length represents interaction strength (i.e. the number of normalised Hi-C links). Putative plasmids with AMR are highlighted yellow.

Lastly, we estimated phylogenetic signal in these super generalists and tested whether AMR presence was associated with phylogenetic signal strength. Phylogenetic signal quantifies the relationship between interaction strength and MAG phylogenetic distance, with high phylogenetic signal indicating that MAGs that are closely related are very likely to have a similar number of interactions with a particular plasmid. Importantly, two main methods are available to quantify phylogenetic signal: the first simply measures autocorrelation between trait differences and phylogenetic distance (measured by Abouheif’s *C*_mean)_; the second applies more complex evolutionary models to test whether distributions match what would be expected if traits coevolved measured by Pagel’s λ (Münkemüller *et al*. 2012). We found that most putative plasmids demonstrated significant phylogenetic signal when measured by both methods (Fig. 5). Contrary to our predictions, putative plasmids carrying AMR had significantly stronger phylogenetic signal than those without when measured by autocorrelation (Fig. 5A) and by an evolutionary model (Fig. 5B), although estimates may be biased by AMR putative plasmids being generally more prevalent than those without AMR (Fig. 5).

**Figure 5.**
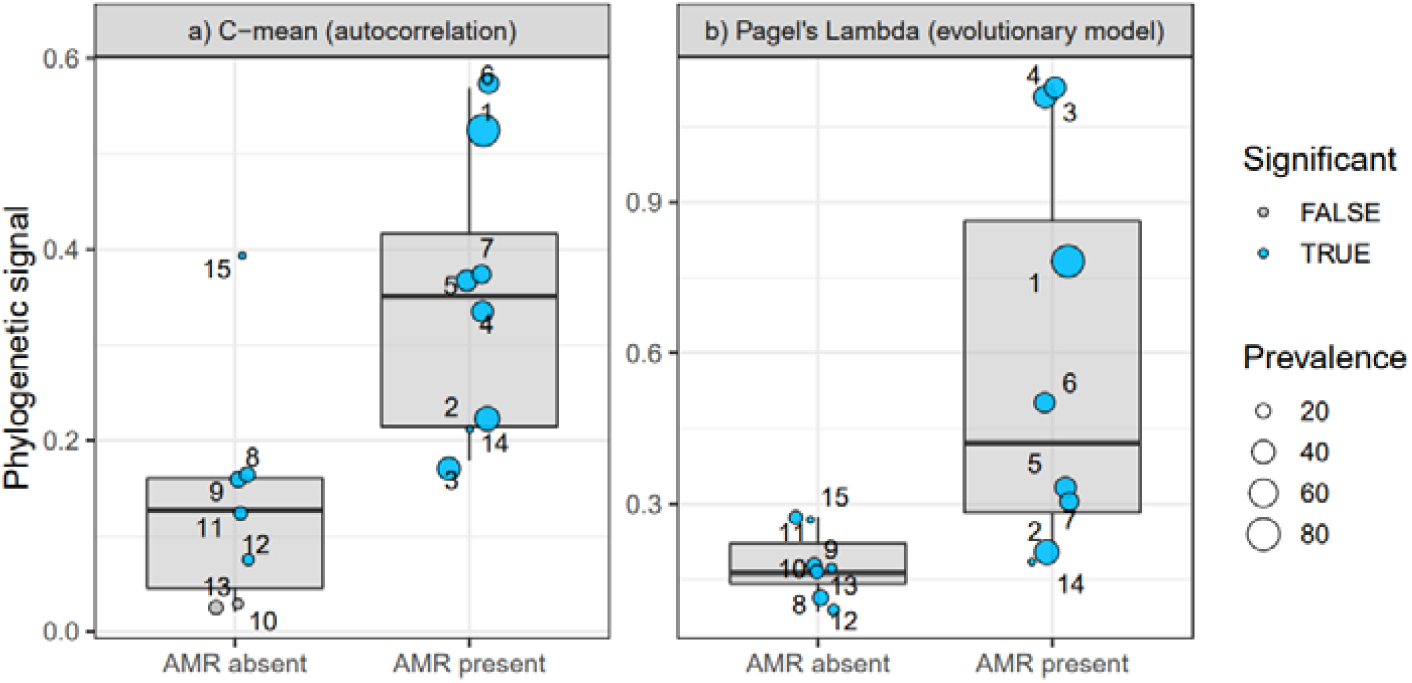
Phylogenetic signal and AMR presence. Phylogenetic signal for the 15 most prevalent putative plasmids separated by AMR presence, applying a) Abouheif’s *C*_mean_, which estimates autocorrelation between trait similarity and phylogenetic distance, and b) Pagel’s λ, which applies a model of trait evolution. Points are labelled by the plasmid ID (corresponding to Fig. 4), sized by their prevalence, and coloured by whether estimates for phylogenetic signal are statistically significant.

## DISCUSSION

The wastewater network analysis demonstrates that while the natural plasmid-host community is dominated by specialist putative plasmids, those carrying AMR genes tend to be more generalist and markedly increase the connectivity of the network. As predicted, the network structure for the AMR plasmid-host subnetwork differed substantially from the non-AMR plasmid network. The AMR plasmid – host network showed a high degree of generalism and nestedness, with an overall high level of connectedness. Extrapolations from ecological network theory (Thebault & Fontaine 2010) and experimental data (Heß *et al*. 2021; Newbury *et al*. in press) suggest that this pattern can be explained by the two different types of plasmids in this network: (1) Costly, more specialised plasmids and (2) beneficial, more generalist plasmids. It is reasonable to think that AMR genes can be directly beneficial in the wastewater environment because they can confer a selective advantage even in the presence of low concentrations of antibiotics and other biocides (Murray *et al*. 2018). Even if not directly beneficial in the current wastewater environment experienced by these organisms, AMR genes will almost certainly have provided a benefit in the environments they originate from, such as hospitals and a community of people consuming various antibiotics. This may have led to interactions that evolved to be more generalised (Guimarães Jr *et al*. 2011; Nuismer *et al*. 2013).).

It is not known precisely why mutualistic networks (e.g., seed dispersal, pollination, symbiosis) often show greater generalism than antagonistic networks (e.g., herbivory and parasitism), with mechanism largely inferred from theory alone. It may be purely short-term ecological consequences, with the greater fitness of mutualistic partners leading to the subsequent spread of mutualists to new species. More generalism may also lead to greater stability in communities dominated by mutualistic interaction (i.e., species are less likely to go extinct, resulting in changes in network properties), while generalism decreases stability of antagonistic communities (Thébault & Fontaine 2010). Theory also suggests that coevolution may drive this pattern under the assumption that trait matching (e.g., attack-defence traits) determines the strength of antagonistic interactions while trait differences (e.g., barriers for transmission) determine mutualistic interactions (Nuismer *et al*. 2013; de Andreazzi *et al*. 2020). While it is impossible to deduce mechanism from our correlational study, recent experiments and theory work using simplified bacteria-plasmid networks demonstrates that short-term growth rate advantages conferred by a beneficial plasmid can result in greater plasmid ecological generalism (Newbury *et al*. in press). Specifically, if a plasmid increases the frequency of its host, the plasmid then has greater opportunities to be transmitted to other host taxa. While the distribution of hosts of a plasmid can be influenced by factors affecting its ability to transfer into a new host, after entry it is primarily the plasmid-encoded replication system and its interaction with host factors that determines the ability of a plasmid to survive in that host (del Solar *et al*. 1996; Toukdarian 2004). This suggests that increased ecological generalism of beneficial plasmids could in turn promote greater evolutionary generalism as a consequence of mutualistic coevolution for plasmid maintenance occurring between plasmids and the multiple hosts they interact with (Harrison & Brockhurst 2012).

We predicted that plasmids with AMR would show lower phylogenetic signal than those without, because mutualists tend to evolve weaker phylogenetic signal than antagonists (Rohr & Bascompte 2014). Yet, we found the opposite: plasmids with AMR genes showed higher phylogenetic signal when measured by both autocorrelation and evolutionary models. This finding may either reflect that AMR plasmids can still be parasitic in some contexts, and therefore require hosts to carefully control plasmid entry and maintenance in a way that might evolutionarily constrain interactions. Another possibility is that AMR plasmids are indeed generally beneficial, yet if most plasmids are parasitic this would still promote the evolution of specific interactions between bacterial and mutualists that act to constrain interactions within phylogenies (Thrall *et al*. 2007). In general, whilst most putative plasmids were highly specialist, super generalist putative plasmids were still largely shared within phyla, and only rarely interacted with bacterial hosts outside of the dominant host phylum. This pattern was irrespective of whether they carried AMR genes or not. This indicates strong barriers to plasmid transmission between phyla (Redondo-Salvo *et al*. 2020), yet not between different classes within phyla. For example, Gamma- and Proteo-bacteria appeared to freely share putative plasmids and did not form separate clusters within networks.

While greater generalism associated with AMR plasmids obviously has important implications for the spread of the specific AMR genes encoded by the plasmids, it is also likely to affect the spread of additional AMR genes, even those not currently under selection. First, a new AMR gene that gets incorporated into a generalist plasmid will have more chance to spread. Second, generalist plasmids, are more likely to acquire additional AMR genes (e.g., by transposition), given the greater diversity of hosts they interact with. Generalist plasmids and in particular generalist plasmids with AMR were mostly associated with Proteobacteria, although plasmid hubs were found across bacterial classes. Generalist AMR plasmids assumed a central role in the overall network by linking Proteobacteria to other classes, although these interactions were relatively rare, and these generalist plasmids may contribute to the spread of AMR gene transfer in general between different classes of bacteria. This might reflect greater selection for AMR in Proteobacteria because many common human pathogens are found in the Proteobacteria.

Our approach advances on the analysis from Stalder et al. (2019) by utilizing novel machine learning methods to identify undescribed plasmid signatures. Whilst this method considerably increases our understanding of how undescribed plasmids contribute to interaction network structure, we assume that connected clusters of sequences represent one plasmid. Yet, it is possible that these clusters in fact represent multiple co-occurring plasmids, or, conversely, that some sequences treated as separate plasmids are in fact part of the same plasmid. Nevertheless, our sensitivity analyses suggest that our results and interpretations are robust to changes to methodology. An additional limitation to our approach is that some shared genes or mobile elements between different plasmids could have amplified the connections of generalist putative plasmids to more hosts. We strived to remove any such genomic elements, such as transposons, AMR, metal resistance, biocide resistance and virulence, yet it is possible that at least some super generalist putative plasmids may be a product of other plasmid accessory genes commonly shared among different plasmids of this bacterial community. Future advances in Hi-C technology paired with long-read sequencing methods will further our ability to distinguish and describe plasmids in natural communities using high throughput sequencing technology. Lastly, this is a correlational study and AMR presence and generalism may also be driven by host taxa. Indeed, proteobacteria are a very ecologically diverse phylum (Woese 1987), so they may be more likely to be associated with promiscuous plasmids carrying genes beneficial in a range of hostile environments. Therefore, to fully understand the ecological and evolutionary dynamics under varying plasmid-host interaction type we need experimental approaches that measure fitness consequences and link those to changes in observed network structures.

By conducting ecological network analyses on a wastewater Hi-C metagenome, we have been able to describe a natural plasmid-host network. The patterns we observe are consistent with theory. First, networks are primarily driven by specialism, consistent with a predominantly parasitic impact of plasmids in the absence of carriage of beneficial accessory genes. Second, greater prevalence of AMR genes – which are often transferred by plasmids – in generalist and abundant plasmids lead to a more connected network. Third, this offers a large potential for sharing of a few generalist plasmids across the network, promoting inter-class HGT and indirect network interactions. Further work is clearly required to determine the generality of our findings and the mechanisms underpinning them. This includes other types of networks, such as bacteria-bacteriophage (Flores *et al*. 2013), where interactions while primarily antagonistic can also be mutualistic (Harrison & Brockhurst 2017; Wendling *et al*. 2021). A closer look at the types of plasmids that cause higher network connectedness would also help understand the drivers of AMR spread in various environments.

## Materials and Methods

### Processing data

#### Generating bacterial MAG data

Hi-C metagenome data from Stalder et al. was assembled into MAGs using an updated algorithm of ProxiMeta™ on April 4th 2021 (Phase Genomics, Inc. 2021). This generated 374 MAGs analyzed in this study. We assigned MAG taxonomy by running MAGs through Phylophlan (Asnicar et al. 2020), which calls MASH for taxonomic assignment, and used taxonomy as a proxy for phylogeny (Table S1).

#### Contig filtering

The raw Hi-C data from one wastewater sample contains over 2.5 million contigs representing DNA fragments from diverse organisms. Because we are interested specifically in links between bacteria and plasmids, we initially filtered out contigs that were 1) identified as transposons, and 2) were not identified as either a MAG, a plasmid, or carrying other AMR, virulence, metal and biocide genes. Transposons and IS elements were identified by performing a homology search with BLASTp on predicted gene from all contigs using an e-value < 0.01 against all known transposase proteins from the databases from IS finder (Siguier *et al*. 2006) available from https://github.com/thanhleviet/ISfinder-sequences/blob/master/README.md and from Tn3 Transposon Finder (Ross *et al*. 2021) available from https://tncentral.proteininformationresource.org/TnFinder.html. Protein-coding genes were predicted from all contigs using prodigal in metagenomic mode using the option ‘-p meta’ available from https://github.com/hyattpd/Prodigal (Hyatt *et al*. 2010). Contigs with gene coding for antimicrobial resistance, virulence factors, metal resistance or resistance to biocides were identified using AMR finder plus (Feldgarden *et al*. 2019), using ‘-n’ and ‘–-plus’ parameters.

#### Identifying putative plasmids

All remaining contigs were run through PlassClass (Pellow *et al*. 2020) using default parameters to distinguish between contigs that were of chromosomal or plasmid origin. This method performs better on short reads < 1000 bp than other machine learning algorithms such as Plasflow (Krawczyk *et al*. 2018). To be conservative, only contigs where 95% of sequence signatures were classified as plasmid signatures. Only one of these was identified as a well-characterised broad-range plasmid (IncP**-**β group). Contigs with gene coding for antimicrobial resistance, virulence factors, metal resistance or resistance to biocides were not treated as belonging to plasmids because, like transposons, they can be shared across multiple plasmids and therefore hinder the identification of unique plasmid signatures. Because the contigs identified as plasmids were mostly constituted by short contigs (the median length was 379 bp) and plasmids are likely to be > 1000 bp long, we reasoned that for a contig to belong to a plasmid it should be found to be consistently connected to at least one other. In order to account for this, we retained only plasmid contigs that were linked to other plasmid contigs at least five times (n = 841 contigs). We then performed a cluster analysis on these plasmid contigs using the Walktrap clustering algorithm using the *igraph::walktrap*.*community* function and with a step length of 4 (Csardi & Nepusz 2006). This clustering step identified 331 plasmid clusters which we treated as putative plasmids.

We conducted several quality checks to assess the reliability of these cluster of contigs we called putative plasmids. We first checked the total length of plasmid contigs. The average total length (4,200 bp) and the general distribution (median = 1,300 bp, min = 500 bp, max. = 78,700 bp) were below typical plasmid length found in natural communities suggesting those clusters of plasmids contigs were part of plasmids but we did not assemble complete plasmids (Dunivin *et al*. 2019). Of these, 39 clusters were found not to associate with any MAGs and were excluded. In addition, we used BLASTn (Megablast against the non-redundant nucleotide database) to manually check the gene content of 132 contigs (out of 841) that were part of putative plasmid clusters that were subsequently found to associate with over 10% of MAGs. Thirty-three contigs were removed at this stage for having genes that could plausibly be associated with transposons or bacterial chromosomes. Lastly, associations characterised by only one Hi-C link were considered unreliable and removed. After this quality filtering, 289 putative plasmid clusters made up of 729 contigs were retained for analysis (Fig. S1a). The remaining putative plasmids we classified as associating with an AMR gene if at least one contig within the cluster was connected to an AMR contig at least twice (Fig. S1b). Hi-C link counts were then normalised Hi-C by both MAG abundance and by putative plasmid size.

#### Analysis

An adjacency matrix was generated from the processed Hi-C association data, and data was handled using the packages phyloseq and igraph. Networks were visualized using the ggnetwork (Briatte 2020) using graphopt layout. Network statistics for the five major host classes were generated with the *bipartite::networklevel* function (Dormann *et al*. 2009). Phylogenetic trees and their attributes were visualized with the *ggtree* package (Yu *et al*. 2017). Phylogenetic signal was estimated using the *phylosignal::phylosignal* function (Keck *et al*. 2016), applying two different measures: Abouheif’s C_mean_, which calculates autocorrelation between phylogenetic distance and trait distributions, and Pagel’s λ, which uses a Brownian motion (BM) model of trait evolution. We chose these two metrics because they perform the best out of a number of metrics available, and are also insensitive to branch length (Münkemüller *et al*. 2012), therefore are appropriate to use on our phylogenetic tree based on taxonomy. To test whether plasmids with and without AMR genes differ in their phylogenetic signal, be performed a t-test.

The Rmarkdown report is available at https://github.com/Riselya/Plasmid-networks. Sequencing data are available in FASTQ format at SRA accession PRJNA506462. Processed data and scripts for linking contigs to genome clusters using Hi-C data are available at https://osf.io/ezb8j/.

## Acknowledgments

We thank Mike Brockhurst for the discussion about the manuscript and Suzanne Kay for help with the data analysis. This research is funded by NERC, grant number NE/S000771/1. BIS is supported by a Royal Commission for the Exhibition of 1851 Research Fellowship. TS is in part supported by the program Understanding Antimicrobial Resistance grant no 2018-67017-27630 / project accession no 1015171 from the USDA National Institute of Food and Agriculture.

## Author Contributions

DS, AB and AR conceived and designed the study. AR, TS, and BIS analyzed the data. DS and AR wrote the first manuscript draft and all authors contributed.

## Supplementary material

**Figure S1.**
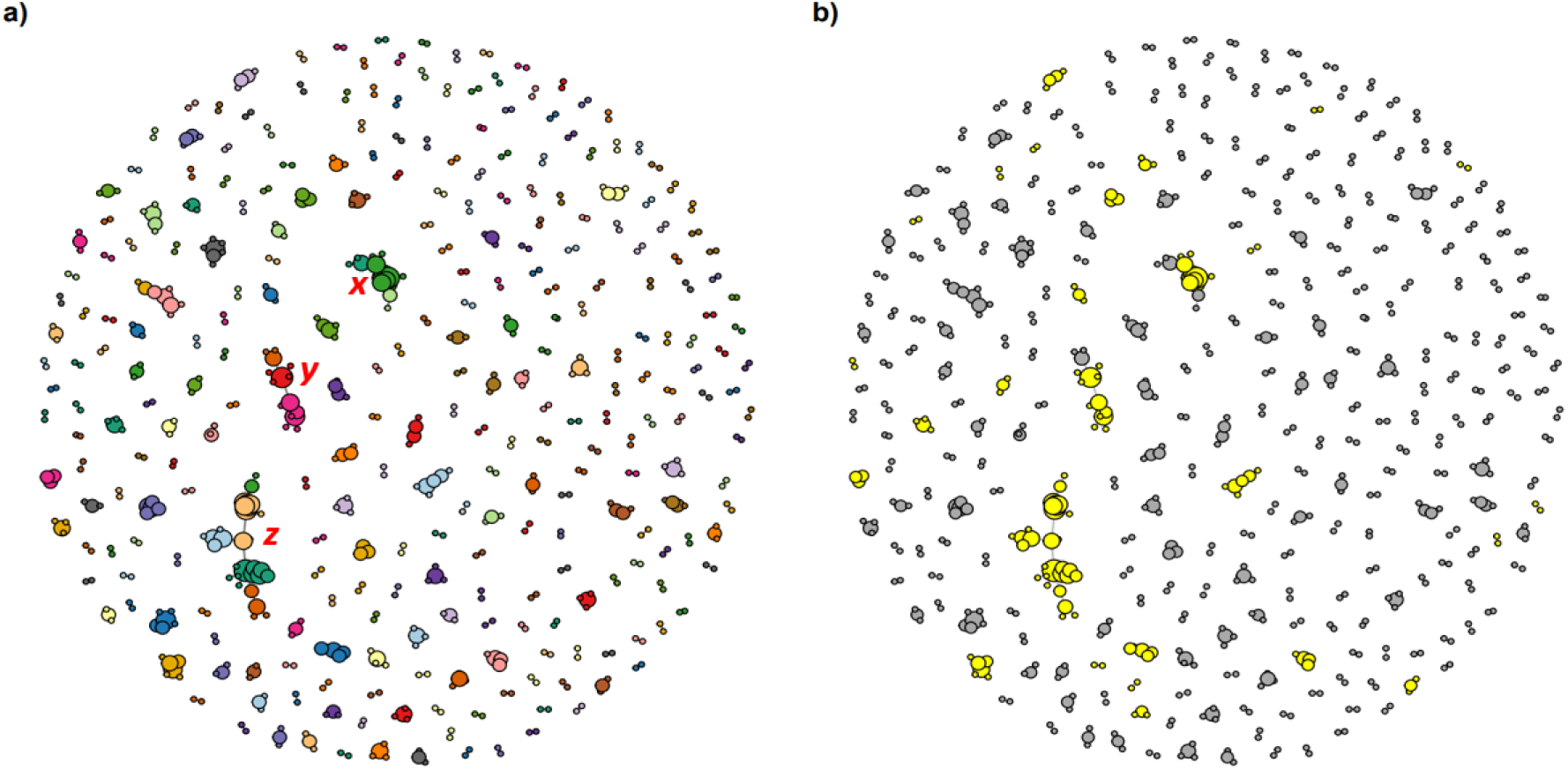
Putative plasmid clusters (n = 289 clusters built from 729 contigs) that were retained for analysis. Each node represents a contig identified as having a 95% probability of belonging to a plasmid by PlassClass, coloured by a) its cluster membership based on the walktrap method, and b) whether it is associated with AMR (yellow). Only associations based on at least 5 Hi-C links are included.

## References

1. Acman, M., van Dorp, L., Santini, J.M. & Balloux, F. (2020). Large-scale network analysis captures biological features of bacterial plasmids. Nature Communications, 11, 2452.

2. Asnicar, F., Thomas, A.M., Beghini, F., Mengoni, C., Manara, S., Manghi, P. et al. (2020). Precise phylogenetic analysis of microbial isolates and genomes from metagenomes using PhyloPhlAn 3.0. Nature Communications, 11, 2500.

3. Bennett, P.M. (2008). Plasmid encoded antibiotic resistance: acquisition and transfer of antibiotic resistance genes in bacteria. British Journal of Pharmacology, 153, S347–S357.

4. Briatte, F. (2020). ggnetwork: geometries to plot networks with ‘ggplot2’. R package version 0.5. 8. 5.

5. Coyte, K.Z., Schluter, J. & Foster, K.R. (2015). The ecology of the microbiome: Networks, competition, and stability. Science, 350, 663–666.

6. Csardi, G. & Nepusz, T. (2006). The igraph software package for complex network research. InterJournal, complex systems, 1695, 1–9.

7. Dang, B., Mao, D., Xu, Y. & Luo, Y. (2017). Conjugative multi-resistant plasmids in Haihe River and their impacts on the abundance and spatial distribution of antibiotic resistance genes. Water Research, 111, 81–91.

8. de Andreazzi, C.S., Astegiano, J. & Guimarães Jr, P.R. (2020). Coevolution by different functional mechanisms modulates the structure and dynamics of antagonistic and mutualistic networks. Oikos, 129, 224–237.

9. del Solar, G., Alonso, J.C., Espinosa, M. & Díaz-Orejas, R. (1996). Broad-host-range plasmid replication: an open question. Molecular microbiology, 21, 661–666.

10. Dimitriu, T., Matthews, A.C. & Buckling, A. (2021). Increased copy number couples the evolution of plasmid horizontal transmission and plasmid-encoded antibiotic resistance. Proceedings of the National Academy of Sciences, 118, e2107818118.

11. Dormann, C.F., Fründ, J., Blüthgen, N. & Gruber, B. (2009). Indices, graphs and null models: analyzing bipartite ecological networks. The Open Ecology Journal, 2.

12. Dunivin, T.K., Choi, J., Howe, A., Shade, A. & Sharpton, T.J. (2019). RefSoil+: a Reference Database for Genes and Traits of Soil Plasmids. mSystems, 4, e00349–00318.

13. Feldgarden, M., Brover, V., Haft, D.H., Prasad, A.B., Slotta, D.J., Tolstoy, I. et al. (2019). Validating the AMRFinder Tool and Resistance Gene Database by Using Antimicrobial Resistance Genotype-Phenotype Correlations in a Collection of Isolates. Antimicrobial Agents and Chemotherapy, 63, e00483–00419.

14. Flores, C.O., Valverde, S. & Weitz, J.S. (2013). Multi-scale structure and geographic drivers of cross-infection within marine bacteria and phages. The ISME Journal, 7, 520–532.

15. Fontaine, C., Thébault, E. & Dajoz, I. (2009). Are insect pollinators more generalist than insect herbivores? Proceedings of the Royal Society B: Biological Sciences, 276, 3027–3033.

16. González-Salazar, C. & Stephens, C.R. (2012). Constructing Ecological Networks: A Tool to Infer Risk of Transmission and Dispersal of Leishmaniasis. Zoonoses and Public Health, 59, 179–193.

17. Guimarães Jr, P.R., Jordano, P. & Thompson, J.N. (2011). Evolution and coevolution in mutualistic networks. Ecology Letters, 14, 877–885.

18. Guimarães, P.R., Pires, M.M., Jordano, P., Bascompte, J. & Thompson, J.N. (2017). Indirect effects drive coevolution in mutualistic networks. Nature, 550, 511–514.

19. Harrison, E. & Brockhurst, M.A. (2012). Plasmid-mediated horizontal gene transfer is a coevolutionary process. Trends in Microbiology, 20, 262–267.

20. Harrison, E. & Brockhurst, M.A. (2017). Ecological and Evolutionary Benefits of Temperate Phage: What Does or Doesn’t Kill You Makes You Stronger. BioEssays, 39, 1700112.

21. Heß, S., Kneis, D., Virta, M. & Hiltunen, T. (2021). The spread of the plasmid RP4 in a synthetic bacterial community is dependent on the particular donor strain. FEMS Microbiology Ecology, 97.

22. Hyatt, D., Chen, G.-L., LoCascio, P.F., Land, M.L., Larimer, F.W. & Hauser, L.J. (2010). Prodigal: prokaryotic gene recognition and translation initiation site identification. BMC Bioinformatics, 11, 119.

23. Jensen, R.B. & Gerdes, K. (1995). Programmed cell death in bacteria: proteic plasmid stabilization systems. Molecular Microbiology, 17, 205–210.

24. Kaiser-Bunbury, C.N., Mougal, J., Whittington, A.E., Valentin, T., Gabriel, R., Olesen, J.M. et al. (2017). Ecosystem restoration strengthens pollination network resilience and function. Nature, 542, 223–227.

25. Karkman, A., Do, T.T., Walsh, F. & Virta, M.P. (2018). Antibiotic-resistance genes in waste water. Trends in microbiology, 26, 220–228.

26. Keck, F., Rimet, F., Bouchez, A. & Franc, A. (2016). phylosignal: an R package to measure, test, and explore the phylogenetic signal. Ecology and Evolution, 6, 2774–2780.

27. Kent, A.G., Vill, A.C., Shi, Q., Satlin, M.J. & Brito, I.L. (2020). Widespread transfer of mobile antibiotic resistance genes within individual gut microbiomes revealed through bacterial Hi-C. Nature Communications, 11, 4379.

28. Klümper, U., Riber, L., Dechesne, A., Sannazzarro, A., Hansen, L.H., Sørensen, S.J. et al. (2015). Broad host range plasmids can invade an unexpectedly diverse fraction of a soil bacterial community. The ISME Journal, 9, 934–945.

29. Krawczyk, P.S., Lipinski, L. & Dziembowski, A. (2018). PlasFlow: predicting plasmid sequences in metagenomic data using genome signatures. Nucleic Acids Research, 46, e35–e35.

30. Lili, L.N., Britton, N.F. & Feil, E.J. (2007). The Persistence of Parasitic Plasmids. Genetics, 177, 399–405.

31. Martínez, J.L. (2008). Antibiotics and Antibiotic Resistance Genes in Natural Environments. Science, 321, 365–367.

32. Montesinos-Navarro, A., Hiraldo, F., Tella, J.L. & Blanco, G. (2017). Network structure embracing mutualism–antagonism continuums increases community robustness. Nature Ecology & Evolution, 1, 1661–1669.

33. Münkemüller, T., Lavergne, S., Bzeznik, B., Dray, S., Jombart, T., Schiffers, K. et al. (2012). How to measure and test phylogenetic signal. Methods in Ecology and Evolution, 3, 743–756.

34. Murray, A.K., Zhang, L., Yin, X., Zhang, T., Buckling, A., Snape, J. et al. (2018). Novel Insights into Selection for Antibiotic Resistance in Complex Microbial Communities. mBio, 9, e00969–00918.

35. Newbury, A., Dawson, B., Klümper, U., Hesse, E., Castledine, M., Fontaine, F. et al. (in press). Fitness effects of plasmids shape the structure of bacteria-plasmid interaction networks. Proceedings of the National Academy of Sciences.

36. Nuismer, S.L., Jordano, P. & Bascompte, J. (2013). Coevolution and the architecture of mutualistic networks Evolution, 67, 338–354.

37. Pellow, D., Mizrahi, I. & Shamir, R. (2020). PlasClass improves plasmid sequence classification. PLoS computational biology, 16, e1007781.

38. Redondo-Salvo, S., Fernández-López, R., Ruiz, R., Vielva, L., de Toro, M., Rocha, E.P. et al. (2020). Pathways for horizontal gene transfer in bacteria revealed by a global map of their plasmids. Nature communications, 11, 1–13.

39. Rohr, R.P. & Bascompte, J. (2014). Components of Phylogenetic Signal in Antagonistic and Mutualistic Networks. The American Naturalist, 184, 556–564.

40. Ross, K., Varani, A.M., Snesrud, E., Huang, H., Alvarenga, D.O., Zhang, J. et al. (2021). TnCentral: a Prokaryotic Transposable Element Database and Web Portal for Transposon Analysis. mBio, 12, e02060–02021.

41. San Millan, A. (2018). Evolution of Plasmid-Mediated Antibiotic Resistance in the Clinical Context. Trends in Microbiology, 26, 978–985.

42. Segar, S.T., Fayle, T.M., Srivastava, D.S., Lewinsohn, T.M., Lewis, O.T., Novotny, V. et al. (2020). The role of evolution in shaping ecological networks. Trends in Ecology & Evolution, 35, 454–466.

43. Siguier, P., Pérochon, J., Lestrade, L., Mahillon, J. & Chandler, M. (2006). ISfinder: the reference centre for bacterial insertion sequences. Nucleic acids research, 34, D32–D36.

44. Stalder, T., Press, M.O., Sullivan, S., Liachko, I. & Top, E.M. (2019). Linking the resistome and plasmidome to the microbiome. The ISME Journal, 13, 2437–2446.

45. Suzuki, H., Yano, H., Brown, C.J. & Top, E.M. (2010). Predicting Plasmid Promiscuity Based on Genomic Signature. Journal of Bacteriology, 192, 6045–6055.

46. Thébault, E. & Fontaine, C. (2010). Stability of Ecological Communities and the Architecture of Mutualistic and Trophic Networks. Science, 329, 853–856.

47. Thrall, P.H., Hochberg, M.E., Burdon, J.J. & Bever, J.D. (2007). Coevolution of symbiotic mutualists and parasites in a community context. Trends in Ecology & Evolution, 22, 120–126.

48. Toukdarian, A. (2004). Plasmid strategies for broad-host-range replication in Gram-negative bacteria. Plasmid biology, 257–270.

49. Veron, S., Fontaine, C., Dubos, N., Clergeau, P. & Pavoine, S. (2018). Predicting the impacts of co-extinctions on phylogenetic diversity in mutualistic networks. Biological Conservation, 219, 161–171.

50. Wang, J., Gao, Y. & Zhao, F. (2016). Phage–bacteria interaction network in human oral microbiome. Environmental Microbiology, 18, 2143–2158.

51. Weitz, J.S., Poisot, T., Meyer, J.R., Flores, C.O., Valverde, S., Sullivan, M.B. et al. (2013). Phage–bacteria infection networks. Trends in Microbiology, 21, 82–91.

52. Wendling, C.C., Refardt, D. & Hall, A.R. (2021). Fitness benefits to bacteria of carrying prophages and prophage-encoded antibiotic-resistance genes peak in different environments. Evolution, 75, 515–528.

53. Woese, C.R. (1987). Bacterial evolution. Microbiological reviews, 51, 221–271.

54. Yaffe, E. & Relman, D.A. (2020). Tracking microbial evolution in the human gut using Hi-C reveals extensive horizontal gene transfer, persistence and adaptation. Nature Microbiology, 5, 343–353.

55. Yu, G., Smith, D.K., Zhu, H., Guan, Y. & Lam, T.T.Y. (2017). ggtree: an R package for visualization and annotation of phylogenetic trees with their covariates and other associated data. Methods in Ecology and Evolution, 8, 28–36.

